# Escalation of alcohol intake is associated with regionally decreased insular cortex activity but not associated with changes in taste quality

**DOI:** 10.1101/2022.10.06.511140

**Authors:** A Mukherjee, MS Paladino, SL McSain, EA Gilles-Thomas, DD Lichte, RD Camadine, S Willock, K Sontate, SC Honeycutt, GC Loney

## Abstract

**Background:** Intermittent access to ethanol (EtOH) drives persistent escalation of intake and rapid transition from moderate to compulsive-like drinking. Intermittent EtOH drinking may facilitate escalation in part by altering aversion-sensitive neural substrates, such as the insular cortex (IC), thus driving greater approach toward stimuli previously treated as aversive.

**Methods:** We conducted a series of experiments in rats to examine behavioral and neural responses associated with escalation of EtOH intake. First, taste reactivity analyses quantified the degree that intermittent brief-access ethanol exposure (BAEE) alters sensitivity to the aversive properties of EtOH. Next, we determined whether pharmacological IC inhibition facilitated EtOH escalation. Finally, given that IC is primary gustatory cortex, we employed psychophysical paradigms to assess whether escalation of EtOH intake induced changes in EtOH taste. These paradigms measured changes in sensitivity to the intensity of EtOH taste and whether escalation shifts the salient taste quality of EtOH by measuring the degree that the taste of EtOH generalized to a sucrose-like (‘sweet’) or quinine-like (‘bitter’) percept.

**Results:** We found a near complete loss of aversive oromotor responses in EtOH-exposed relative to -naïve rats. Additionally, we observed significantly reduced expression of EtOH-induced c-*Fos* expression in the posterior IC in exposed rats relative to naïve rats. Inhibition of the IC resulted in a modest, but statistically reliable increase in acceptance of higher EtOH concentrations in naïve rats. Finally, we found no evidence of changes in the psychophysical assessment of the taste of EtOH in exposed, relative to naïve, rats.

**Conclusions:** Our results demonstrate that neural activity within the IC adapts following escalation of EtOH intake in a manner that correlates with reduced sensitivity to the aversive hedonic properties of EtOH. These data further establish that IC may be driving exposure-induced escalations in EtOH intake and directly contributing to development of compulsive-like EtOH drinking.

## 1.0 Introduction

A key feature of alcohol use disorders (AUDs) is the transition from recreational to escalated use that persists despite adverse consequences. A similar phenomenon exists in rodent models, as repeated, intermittent access to ethanol (EtOH) drives escalation of voluntary intake of EtOH, particularly at higher concentrations which are often treated as aversive in naïve rats (Carnicella et al., 2014, Simms et al., 2008, Rosenwasser et al., 2013, Lundqvist et al., 1994). Intermittent-EtOH access and/or presentation of EtOH-predictive cues can sufficiently escalate intake above that observed in continuous-access paradigms, mimicking moderate to excessive EtOH drinking behaviors in humans (Carnicella et al., 2014) and this occurs whether EtOH is presented intermittently across days (Simms et al., 2008), or within session (Tomie et al., 2006, Loney and Meyer, 2018). We previously demonstrated that intermittent brief-access (≤10s) to multiple EtOH concentrations in 30-min sessions (brief-access EtOH exposure; BAEE) is sufficient to induce rapid escalation of intake in rats to levels typically observed following intermittent 24-h two-bottle choice tests (Loney and Meyer, 2018). Intermittent-EtOH intake often precedes the development of compulsive-like drinking behavior in humans (Sanchez-Marin et al., 2017) which is often conceptualized as reductions in sensitivity to the aversive properties of EtOH consumption and continued drinking despite deleterious consequences (Hopf et al., 2010, Hopf and Lesscher, 2014, Chen et al., 2015). While the neural mechanisms underlying escalation have yet to be fully characterized, there is evidence to suggest that a key driver of augmented consumption is decreased sensitivity to the aversive properties of EtOH and increasing evidence supports a role for insular cortex (IC) in mediating sensitivity to the aversive consequences of psychoactive substances (Gehrlach et al., 2019, Contreras et al., 2007, Venniro et al., 2017, Seif et al., 2013, Campbell et al., 2019, Campbell and Lawrence, 2021, Geddes et al., 2008, Mackey et al., 1986). IC is extensively and reciprocally connected with many subcortical regions involved in drug-seeking and taking (Naqvi and Bechara, 2010) including receiving dopaminergic projections from the ventral tegmental area (VTA) and sending projections to the ventral striatum, which may modulate the reinforcing efficacy and interoceptive effects of psychoactive compounds (Zito et al., 1988, Naqvi and Bechara, 2010, Ohara et al., 2003). IC signaling is necessary for processing aversive somatosensory and interoceptive information, which may protect the user from EtOH overconsumption by signaling the salience of its aversive interoceptive properties (Schier et al., 2014, Berret et al., 2019).

In addition to the critical role IC plays in processing drug-related extraneous cues, it is also the location of primary gustatory cortex thus being an essential brain area for taste processing and modulation, with both IC afferents and efferents involved in the mediation of taste learning (Cubero et al., 1999, Roman and Reilly, 2007, Schier et al., 2014, Yiannakas and Rosenblum, 2017, Abe et al., 2020, Jung et al., 2022). Lesions of IC have been shown to disrupt acquisition of conditioned taste aversion (CTA) and impair taste sensitivity to quinine and potassium chloride, but not sucrose, suggesting a potential biased role in relaying information for aversive taste stimuli (Bales et al., 2015). Additionally, differences in the degree of activation of various subregions within IC, represented by *Fos*-like immunoreactivity (FLI), is predictive of the strength of aversive and rejection responses to intraoral infusions of taste stimuli (King, 2018). Given the overlap of IC with gustatory cortex, one hypothesis for escalation of EtOH intake following intermittent access is exposure-induced alterations in EtOH-taste perception mediated, at least in part, by altered IC neural activity. Here, we aimed to elucidate the mechanisms and parameters by which BAEE robustly increases voluntary consumption of higher concentrations of EtOH. First, we investigated whether BAEE alters the hedonic taste response to EtOH by quantifying appetitive and aversive taste responses to EtOH following prior experience with BAEE. Additionally, we quantified the FLI generated by intraoral EtOH across three levels of IC (anterior, mid, and posterior). Next, we pharmacologically inhibited IC of EtOH-naïve rats immediately prior to BAEE to investigate whether reduced activation in IC results in enhanced initial acceptance of higher EtOH concentrations. Finally, we assessed whether the perceived taste intensity, or quality, of EtOH changed following escalation under the BAEE model. We tested BAEE-exposed and -naïve rats in an EtOH-taste detection paradigm to assess differences in the perceived taste intensity of EtOH and examined whether EtOH exposure alters the perceived salient taste quality of various EtOH concentrations. These findings contribute to the literature on the role played by IC in escalation of EtOH intake and exposure-induced changes in the hedonic responding to EtOH which likely play an integral role in transitioning from recreational to compulsive-like drinking.

## 2.0 Methods

### 2.1 Animals

Adult male and female Wistar rats (n=68, Envigo; Indianapolis, IN) were singly housed in polycarbonate cages in temperature and humidity-controlled rooms maintained on a 12:12h reverse light cycle. Animals were provided with rodent chow (Teklad 2018) and *ad libitum* water in their home cages unless otherwise noted. Following habituation to the facility, all animals were handled for three days prior to the start of any experiments. Experimental procedures were approved by the University at Buffalo Institutional Animal Care and Use Committee.

### 2.2 Taste Stimuli

Taste stimuli were prepared daily in water and consisted of EtOH (Decon Labs) diluted from 200-proof stock solution, quinine hydrochloride (QHCl; Sigma Aldrich), and sucrose (Wegmans). All stimuli were presented at room temperature.

### 2.3. Apparatus

#### Davis Rig

Brief-access EtOH exposure (BAEE) was conducted in a lickometer system called a Davis Rig (Med-Associates) that our lab, and others, have previously used for EtOH exposure in rodent models (Youngentob and Glendinning, 2009, Brasser et al., 2012, Loney and Meyer, 2018). This device (Smith, 2001) consists of an automated table containing 16 fluid reservoirs connected to sipper tubes and a polycarbonate cage with wire-bottom floor. Access to the lickometer-coupled sipper tubes is occluded by a computer-controlled shutter. Each trial, the rat receives brief access to a single spout in the center of the cage only when the shutter is open. Licks on the spout are measured by computer software and stored for offline analyses.

#### Gustometer

Psychophysical testing was conducted in custom-built operant chambers termed gustometers. The gustometer is a precision-designed stimulus delivery system often used to assess taste sensitivity in rodents (Geran and Spector, 2000, Loney et al., 2012, Bales et al., 2015, Spector et al., 2015). These devices consist of a modified skinner box with wire-bottom floor housed in sound attenuating chambers. The gustometer sidewall has 3 openings allowing animals access to 3 response balls: the sample ball (center location) is a borosilicate ball spinning on a horizontal axis; the 2 reinforcement balls (flanked to the left and right of the sample ball) remain stationary. Computerized syringe pumps dispense fluid onto the sample ball from fluid reservoirs connected through polyurethane tubing. The sample ball is housed on an automated, retractable swivel arm. Following the sampling phase, this arm is retracted out of reach and the ball is washed with deionized (DI) water, dried with pressurized air, and repositioned. DI water is similarly dispensed to the reinforcement balls from computerized syringe pumps connected through PE tubing. Licks to all stimuli are recorded by force transducers. White noise is played to limit extraneous sounds that may serve as a non-taste cue to direct responding. A stainless-steel barrier is positioned between the housing-chamber and the stimulus delivery system to limit visual cues. To minimize olfactory cues, an exhaust fan is positioned above the sample ball.

### 2.4 Experiment 1: Changes to Hedonic Ethanol Taste Response Following Prior Exposure to Ethanol Under the BAEE Model

#### 2.4.1 Animals

Following habituation, weight-matched animals were divided into two groups: EtOH-exposed (n=10; 5 females) and -naïve (n=10; 5 females).

#### 2.4.2 Brief Access Ethanol Exposure (BAEE)

##### Training

Training in the Davis rig was modeled after previously published procedures (Loney and Meyer, 2018). Briefly, rats were trained to consume water in the rig and acclimated to movement of the shutter and table. Training sessions lasted for 30 minutes and consisted of water as the sole stimulus. Full training methods are available in the Supplementary Materials online.

##### Testing

Test sessions were identical to the final day of training except that Exposed-animals were presented with a randomized block consisting of six concentrations of EtOH (1.25, 2.5, 5.0, 10, 20 & 40%) and water. Animals were able to initiate as many trials as possible during the 30-minute session. Testing was conducted intermittently on Monday, Wednesday, and Friday of a given week, allowing ∼48 hours between test sessions. Immediately following testing, animals received *ad libitum* food and water overnight in their home cage.

#### 2.4.3 Surgical Procedure

Rats underwent fixation of bilateral intraoral catheters as described previously (Eckel and Ossenkopp, 1995, Loney and Eckel, 2021). Briefly, under isoflurane anesthesia (2-4%), a 7-gauge needle was inserted subcutaneously at the scapular region and guided subcutaneously emerging through the oral mucosa just lateral to the maxillary molar. PE tubing (BD Intramedic) was inserted through the needle, and threaded through a Teflon washer held against the oral mucosa. This procedure was repeated identically on the other side. Animals received postoperative carprofen (s.c, 5mg/kg) and were maintained on wet mash (Teklad 2018) for two days following catheterization. Both catheters were flushed daily with ∼1 ml saline. Behavioral testing was performed following one week of post-surgical recovery.

#### 2.4.4 Oral Infusions

Rats were habituated to the test chamber across three 30-min habituation sessions. These sessions were modeled after previous literature (Loney and Eckel, 2021). On day 1, rats were placed into the chamber and left undisturbed. On day 2, rats were connected to the infusion pump (Cole-Palmer) through tubing connected to an adapter on the end of the oral catheter. Rats were left undisturbed in the testing chamber and did not receive delivery of any solution. On day 3, rats were habituated to intraoral water infusions. The pump infused 5ml of water over ∼20-minutes (0.233ml/min). On the test day, rats received an infusion of 20% EtOH. Immediately after commencing EtOH infusions, rats were videotaped with a digital camera (GoPro) positioned directly below the plexiglass chamber providing direct visualization of all oromotor responses.

#### 2.4.5 Brain Histology

45 min following oral infusions, rats were given an overdose of sodium pentobarbital (150 mg/kg) and transcardially perfused with chilled saline and phosphate-buffered 4% paraformaldehyde (PFA). Brains were removed and post-fixed in PFA overnight at 4°C. Following fixation, brains were cryoprotected with 30% sucrose in 0.1 M phosphate-buffered saline (PBS) plus 0.1% sodium azide. Brains were coronally sectioned at 40μm using a freezing stage microtome (Hacker Instruments) and stored in 0.1M PBS/0.1% azide.

Immunohistochemical staining was conducted by rinsing free-floating brain sections in multiple 0.1 M PBS washes, followed by 0.3% triton-X in 3% normal goat serum (NGS) blocking solution. Next, sections were incubated in primary c-*Fos* antibody (1:1000, ABE457, EMD Millipore) overnight at 4°C.Tissue sections were then washed in a series of 0.1 M PBS washes, blocked with NGS, then incubated in secondary antibody (1:200, CF-488A, Biotium) for 2 hours at room temperature. *Fos*-stained sections were mounted on slides and imaged with a fluorescent microscope equipped with a digital camera (Echo Revolve).

#### 2.4.6 Taste Reactivity Analysis

The first minute of the video was hand-scored and analyzed for taste reactivity responses (Loney and Eckel, 2021, Grill and Norgren, 1978). Oromotor and somatic movements were classified as either appetitive (rhythmic mouth movements, RMM; paw licks, PL; and tongue protrusions) or aversive (gapes and paw flails, PF). Videos were analyzed by an experimenter blinded to exposure condition by playing back the clip at 1/5th speed. Three animals were excluded from analyses due to obscuring of their oral cavity preventing accurate quantification of oromotor movements.

#### 2.4.7 *Fos*-like Immunoreactivity Quantification

Three separate images per rat of the IC were obtained at 10x representing the following approximate anterior-posterior levels: anterior (AP: +2.28), middle (AP: +0.96) and posterior (AP: −0.36). Images and quantification were focused approximately on the dorsal agranular region of the IC (King 2018). Two experimenters blinded to experimental condition manually counted c-*Fos* positive cells in each of the three images. Three animals were excluded from the total quantification due to inadequate fixation and immunohistochemical staining.

### 2.5 Experiment 2: Pharmacological Inhibition of the Insular Cortex Prior to BAEE

#### 2.5.1 Animals

Following habituation, weight-matched animals were divided into experimental and control groups. One group was assigned to receive baclofen-muscimol infusions (BM; n=8; 4 females) and the other phosphate-buffered saline (PBS; n=11; 6 females).

#### 2.5.2 Surgical Procedure

Rats underwent surgery to implant bilateral cannulae (Plastics One) targeting the IC (A/P: +1.0mm, M/L: +/−5.5mm, D/V: −4.4mm) under inhaled anesthesia (2-4% isoflurane). Cannulae were fixed to the skull using dental acrylic (Keystone Industries) and stainless-steel screws. Carprofen (5mg/kg, s.c.) was administered for two days following surgical procedures. Animals recovered for 1 week following surgery prior to behavioral testing.

#### 2.5.3 Brief Access Ethanol Exposure (BAEE) Following Pretreatment with Intra-IC GABA Agonist Cocktail Prior to BAEE

The GABA agonist cocktail (50ng baclofen + 50ng muscimol; 1μl), or PBS, was bilaterally infused with an infusion pump at a rate of 0.5ul/min. Injectors were left in place for an additional minute to allow for adequate drug diffusion. Immediately following infusions, rats were placed into the Davis rig for testing. Training and testing procedures were identical to experiment 1. Rats were presented with a randomized block consisting of six concentrations of ethanol (1.25, 2.5, 5.0, 10, 20 & 40%) and water within 30-min sessions on M, W, and F. Animals received six non-consecutive test sessions (2 weeks) of BAEE. Microinfusion of BM or sterile PBS were given prior to the first three test sessions, and sham injections were given for the following three sessions.

#### 2.5.4 Brain Histology

Following testing, animals were sacrificed and transcardially perfused with chilled saline and buffered 4% PFA. Brains were removed and post-fixed in PFA overnight at 4°C. Following fixation, brains were cryoprotected with 30% sucrose in 0.1M PBS. 40 μm sections were mounted on slides and stained with a 0.1% thionin solution (Sigma-Aldrich) to verify cannula placements within the IC.

### 2.6 Experiment 3a: Ethanol Taste Detection Assessment in Naïve- and Exposed-Rats Following Escalation

#### 2.6.1 Animals

Following habituation, weight-matched animals were divided into two groups: exposed (n=6; 3 females) and naïve (n=6; 3 females). On all weekdays, animals were maintained on a 23-hour water deprivation schedule to facilitate motivation to earn water during the daily testing sessions. Water was available *ad libitum* on weekends. Full methods can be found in the Supplementary Materials online.

#### 2.6.2 Brief Access Ethanol Exposure (BAEE)

Training and testing procedures were identical to experiments 1 & 2. For the testing phase, naïve-rats received water from all spouts during Davis rig sessions, whereas exposed-rats were given 30-min access to the six concentrations (1.25-40%) of EtOH or water in the Davis rig.

#### 2.6.3 Gustometer Spout Training

Rats were initially trained to lick a dry sample stimulus delivery ball located in the center of the front of the chamber. Licking was detected by a force transducer and initiated water delivery to the ball. On subsequent days, rats underwent identical training sessions for the right and left reinforcement balls. This familiarized rats with the device and trained them to obtain fluid from each ball. All rats had access to the same stimulus ball on any given day, and training lasted for 3 days, 1 day per ball. All training and testing sessions were 30 min.

#### 2.6.4 Gustometer Side Training

Next, rats were conditioned to associate a given reinforcement ball with either the standard taste stimuli (sucrose and QHCl) or water. A correct response on the associated reinforcement ball was reinforced with water. Side assignments were counterbalanced across animals. Only one standard stimulus (600mM sucrose, 1.0mM QHCl, or DI water) was presented per day and the other stimuli were introduced on alternating days such that training with sucrose or QHCl was followed by training with water.

#### 2.6.5 Gustometer Alternation Training

At this stage, rats were presented with one taste stimulus (1.0mM QHCl or 600mM sucrose) repeatedly until a criterion number of correct responses occurred (each day the criterion was decreased from 6 to 4 then 2). Correct responses did not have to occur sequentially. Once criterion for that given stimulus was met, the rat was presented with water and required to reach the same criterion before the original stimulus was returned. This pattern was repeated throughout the session and the other stimulus was tested on the next day.

#### 2.6.6 Gustometer Discrimination Training Stage (I–II)

For Stage I of discrimination training, the concentration of each standard stimulus was lowered (200mM sucrose and 0.4mM quinine) and only one standard taste stimulus was tested on a given day. Standard stimulus and water trials were randomized. After reaching a set criterion (∼80% correct discrimination of tastant from water trials), animals moved to Stage II. During stage II, two concentrations of quinine (1.0 & 0.4mM) or sucrose (600 & 200mM) were presented in randomized trials with water and only one taste stimulus was tested per day. On this stage of training, it was found that EtOH-naïve and -exposed rats performed worse on 0.4 mM QHCl (57.9 ± 6.0 and 58.0 ± 5.8% correct, respectively) relative to 1.0 mM QHCl (89.8 ± 5.4 and 87.5 ± 5.6%). Therefore, all animals were given five more days of training between water and 0.4 mM QHCl. At the end of this period, performance on 0.4 mM QHCl increased to 87.2 ± 4.3 and 85.0 ± 3.0% correct for EtOH-naïve and -exposed rats, respectively, so all rats advanced to testing.

#### 2.6.7 Testing EtOH Detection Threshold

The testing phase was modeled after Stage II discrimination training, with the exception that, along with water, one EtOH concentration was presented per session (5%, 3.75%, 2.5%, 1.25%, 0.94%, 0.63%, 0.47, 0.4% v/v), instead of the standard stimuli. Each EtOH concentration was tested on two separate days, and performance was averaged across days. To maintain stimulus control, each test day in which a lower concentration of EtOH was presented was followed with a test day using the highest concentration at which rats reliably performed at, or above, 80% correct.

### 2.7 Experiment 3b: Assessment of the Salient Taste Quality of Ethanol in Naïve- and Exposed-Rats Following Escalation

#### 2.7.1. Animals

Male and female rats were weight-matched and assigned to EtOH-exposed (n=9; 5 females) or -naïve groups (n=8; 4 females). As in Experiment 3a, rats were maintained on an ∼23-hour water deprivation schedule and water was provided *ad libitum* on weekends. These animals were experimentally naïve but had previously undergone a sham intercranial infusion surgery four weeks prior to training.

#### 2.7.2 Brief Access Ethanol Exposure (BAEE)

Rats underwent identical EtOH exposure training and testing as previously described above. For the testing phase, naïve animals received water from all spouts during Davis rig sessions, whereas exposed rats were given 30-min access to water and the six concentrations of EtOH in the Davis rig.

#### 2.7.3. Gustometer Spout Training

As in Experiment 3a, rats were given a three-day training to learning to lick a dry sample stimulus delivery ball at the center, right, or left side to elicit water delivery. Training lasted for 3 days, 1 day per ball. The duration of all training and testing sessions was 30 minutes.

#### 2.7.4. Gustometer Side Training

Rats were trained to associate a given reinforcement ball with a standard stimulus (sucrose or QHCl). Rats were presented with one standard stimulus per day, and only the highest concentration was used (600mM sucrose or 1.0mM QHCl). For half of the rats, sucrose was paired with the right reinforcement ball and QHCl with the left while the other half had the opposite assignments. All other conditions for side training were identical to those described in Experiment 3a.

#### 2.7.5. Gustometer Alternation Training

Alternation training consisted of presenting one of two standard stimuli (600 mM sucrose or 1.0 mM QHCl) repeatedly until rats reached a criterion number of correct responses (6, 4, or 2 on training Days 1, 2, and 3, respectively) as in Experiment 3a. Once criterion was reached, the other stimulus was presented and the animal was required to reach the same correct response criterion before returning to the original stimulus. This pattern was repeated for the duration of the 30-min test.

#### 2.7.6. Gustometer Discrimination Training (Stage I-II)

For the first stage of discrimination training, the intermediate concentrations of the standard stimuli (200mM sucrose and 0.3mM QHCl) were presented in randomized blocks. Rats were tasked with making a choice regarding the taste quality of the sample stimulus by responding on one of the two reinforcement spouts. After reaching a performance criterion (∼80% correct for both stimuli), animals were advanced to Stage II of discrimination training, which consisted of presenting all three concentrations of sucrose and QHCl in blocks of six randomized trials. For sucrose, the three concentrations provided were 60, 200, and 600mM, and for QHCl, the concentrations were 0.1, 0.3, and 1mM. Stage II discrimination training parameters served as a training refresher on the Monday sessions during testing.

#### 2.7.7. Taste Discrimination Testing

The testing phase modeled Stage II discrimination training, except that a single EtOH concentrations was introduced on Tuesday–Friday of each week interspersed with sucrose and QHCl trials. Monday of any given week served as a stimulus control day in which only sucrose and QHCl were presented, with concentrations and randomized presentation of stimuli identical to Stage II discrimination training parameters. Two concentrations of EtOH were tested within a given week, and each concentration of EtOH was tested on two days, with performance averaged across the two testing days. The concentrations of EtOH tested consisted of the same concentrations used in the Davis rig (1.25, 2.5, 5.0, 10, 20, 40% v/v).

### 2.8 Data Analyses

Licks from the Davis Rig were converted into lick ratios by dividing the animal’s licks at each EtOH concentration by that same animal’s licks to water to control for individual differences in local lick rate. A lick ratio of 1 indicates the animal took as many licks to a given EtOH concentration as it did water. For these analyses, we averaged the first three test sessions together and compared them to the average of the final three sessions (Loney and Meyer, 2018).

Performance on the EtOH *taste detection paradigm* is presented as the percentage of correct responses representing the average of the performance on both water and EtOH trials across the entire session. A score of 100 indicated that the animals correctly identified each trial while a score of 50 indicates that the animals were responding at chance levels. Curves were fit to these data with a sigmoidal 3-parameter logistical function:

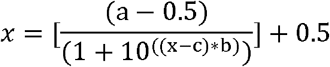

Curve parameters were identified as maximum performance (*a*), slope (*b*) and the concentration at which half-maximal performance was seen (*c*). Variable *x* represents the log-transformed EtOH concentration.

Data from the *taste generalization paradigm* were converted into a Sucrose Generalization Score (Grobe and Spector, 2008; Loney et al., 2012) which represents the degree to which EtOH resembled the taste of sucrose as a function of the animal’s performance on the highest concentrations of the standard sucrose and QHCl trials. The equation used:

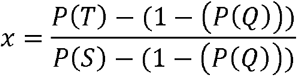

Where P(T) is the proportion of sucrose responses when presented with an EtOH trial, P(Q) is the performance on 1.0mM QHCl and P(S) is the performance on 600mM sucrose.

All data were analyzed with ANOVAs or *t*-tests where appropriate and as identified in the relevant results sections. Data were analyzed with Statistica 12 software and curves were fit with Systat.

## 3.0 Results

### 3.1 Prior BAEE reduces aversive oromotor taste response to EtOH and c-*Fos* expression in posterior IC

Consistent with previous data (Loney and Meyer, 2018), exposure to EtOH within the BAEE paradigm significantly escalated EtOH consumption and increased acceptance of higher concentrations of EtOH. A two-factor mixed ANOVA revealed a significant Test x Concentration interaction (*F*_(5,45)_=11.34, *p*<0.0001). Post-hoc analyses determined that, on average, rats licked significantly more to the 2.5, 5.0, 10, & 20% concentrations following EtOH exposure (Fig 1A). Consistent with this finding, we found a significant escalation in average total EtOH (g/kg/30min) intake from the first (0.3 g/kg) to last (1.01 g/kg) BAEE session (*t*_(10)_ = 6.41, *p*<0.001).

**Figure 1:**
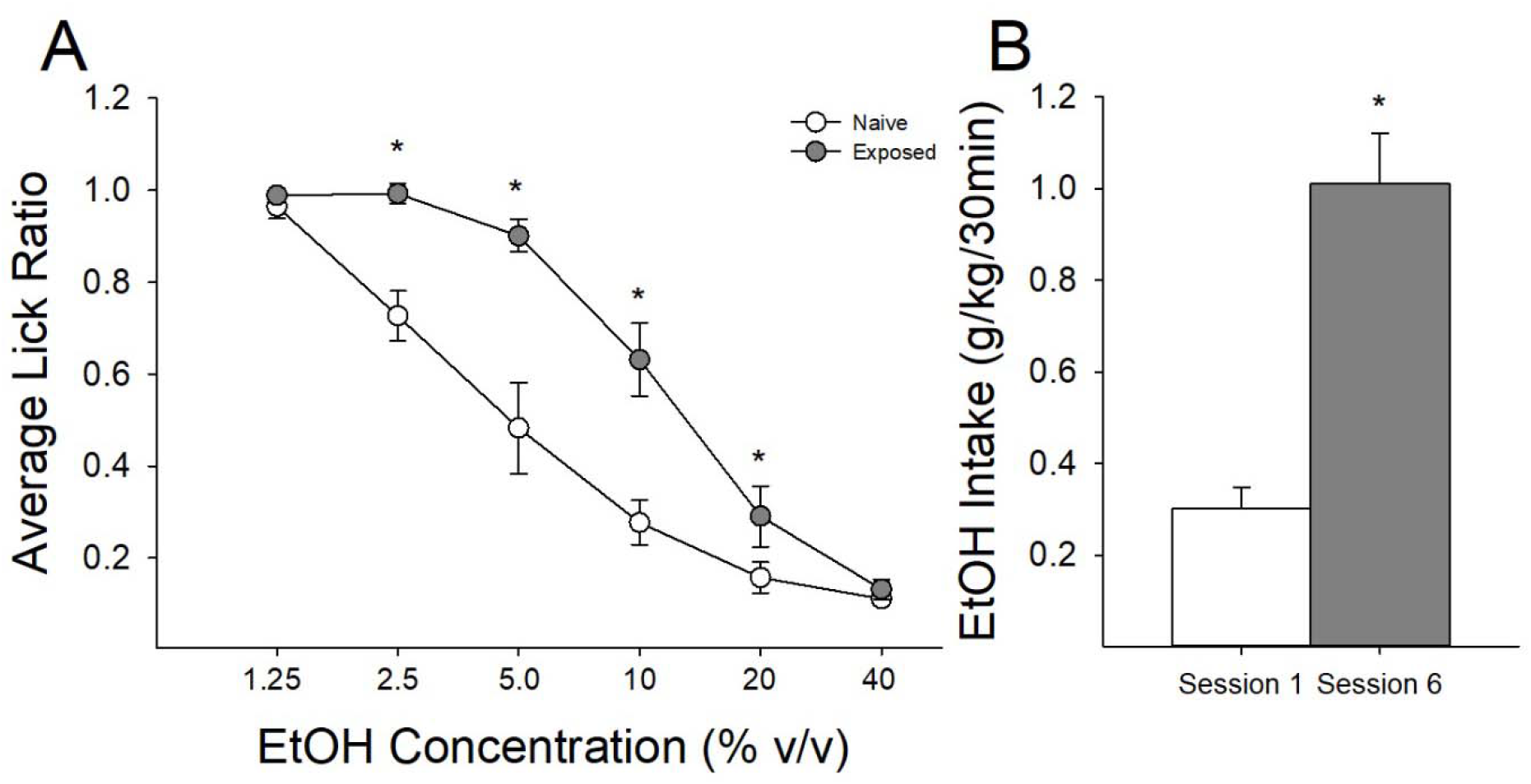
Intermittent brief-access to EtOH results in a significant increase in the acceptance of higher concentrations of EtOH and significantly escalates EtOH consumption. Lick ratios are an individual rat’s licks to a given EtOH concentration compared to that same rat’s licks to water. **(A)** Following six sessions of BAEE rats demonstrate a rightward shift in the acceptance of ETOH resulting in significantly more licks to 2.5%, 5%, 10% and 20% EtOH. **(B)** This rightward shift drives a significant escalation in total EtOH intake (g/kg/30min) from the first session to the sixth BAEE session. * indicate significant group differences (*p*<0.05).

Next, we compared taste reactivity responses to intraoral 20% EtOH between EtOH-exposed and EtOH-naïve rats. Oromotor responses to intraoral EtOH infusions were quantified for the first minute of the taste reactivity video recording. Independent samples t-tests revealed that animals previously exposed to EtOH displayed more appetitive responses elicited by intraoral 20% EtOH infusion (Fig 2A); however, this was not statistically significant (t_(15)_= 1.67; p= 0.114). Conversely, the aversive responses elicited by intraoral 20% EtOH infusion were significantly lower in exposed animals (Fig 2B; *t*_(15)_= 6.41; *p*<0.01), demonstrating that BAEE significantly reduces the aversive responding to concentrated EtOH solutions.

**Figure 2:**
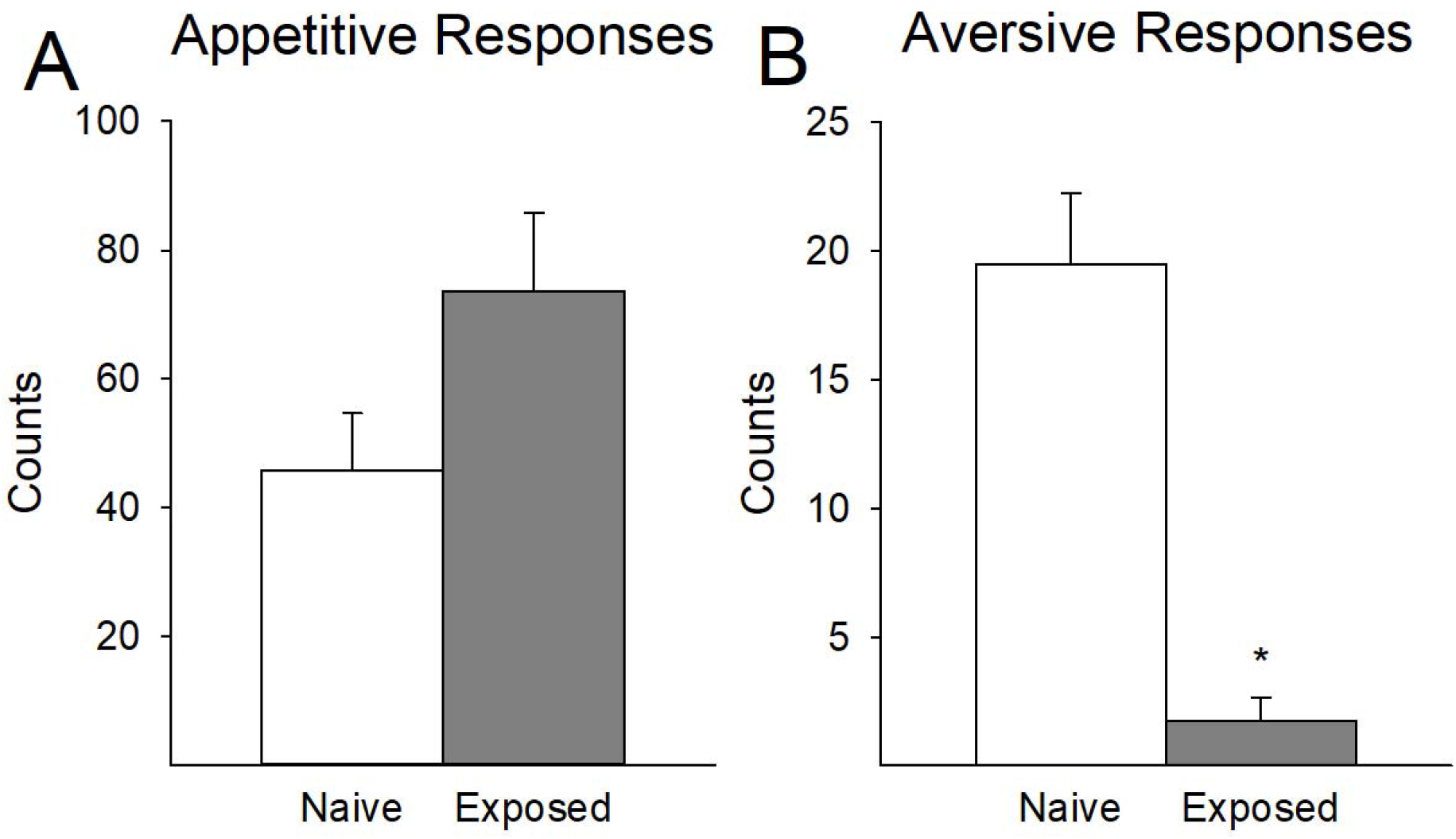
Prior EtOH exposure results in a near elimination of the aversive oromotor responses to orally delivered 20% EtOH. **(A)** Average number of appetitive responses (Rhythmic Mouth Movements, Paw Licks and Tongue Protrusions) was not statistically different between rats previously exposed to EtOH and naïve rats. **(B)** Exposed rats displayed a significant reduction in aversive taste responses (Gapes and Paw Flails) following intraoral administration of 20% EtOH. * indicate significant group differences (*p*<0.05).

Finally, we analyzed the number of FLI-positive cells generated by intraoral 20% EtOH in both EtOH-exposed and -naïve rats. FLI+ cell counts were obtained at three separate anterior-posterior (anterior, mid, posterior) areas of the IC. A two-factor mixed ANOVA revealed a significant IC Area x Condition (Exposed vs. Naïve) interaction (*F*_(2,28)_= 14.25 *p*<0.0001). Post-hoc analyses demonstrated a significant reduction in FLI+ cells in the posterior IC in EtOH-exposed rats relative to naïve rats (Figure 3C). We also ran regression analyses comparing the total FLI+ cells with the number of aversive oromotor responses produced during the taste reactivity analyses following intraoral 20% EtOH in the rats that were included in both analyses. We found a negative correlation between the number of FLI+ cells in the anterior IC and aversive responses, but this effect did not quite reach statistical significance (Fig 4A; *r* = 0.39, *p* = 0.17). There was no discernable relationship between aversive responding and FLI+ markers in the mid IC (Fig 4B; *r* = 0.12, *p* = 0.69). We did find a significant negative correlation between aversive responding and FLI+ cells within the posterior IC (Fig 4C; *r* = 0.71, *p* < 0.01).

**Figure 3:**
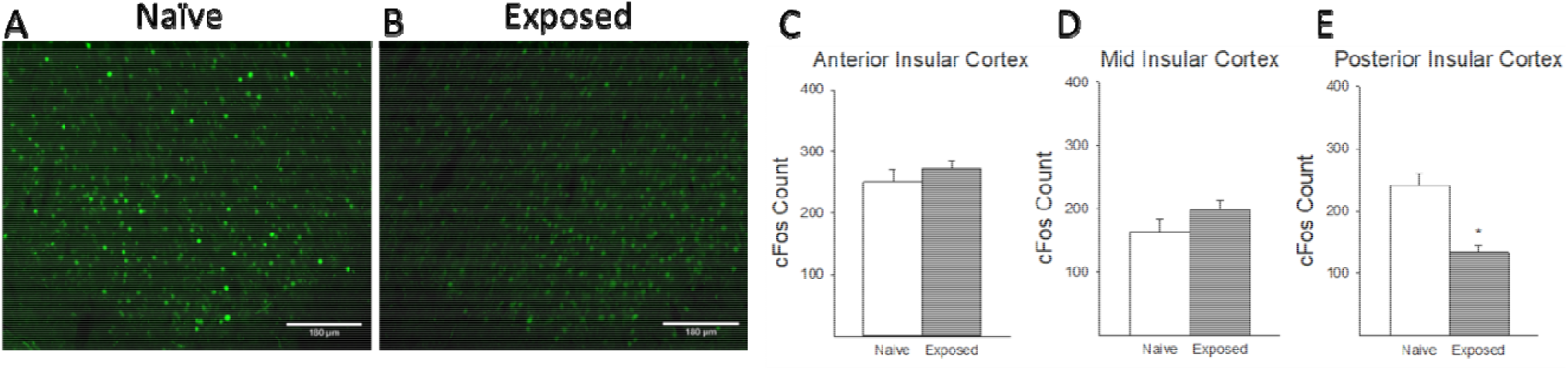
Prior exposure to EtOH results in a significant reduction in *Fos*+ neurons within the posterior insular cortex generated by intraoral delivery of 20% EtOH. **(A & B)**. Representative images of c-*Fos* immunolabelling in the IC in naïve and EtOH-exposed rats. **(C)** There were no differences in the amount of FLI in the anterior or **(D)** mid insular cortex between exposure groups. **(E)** There was a significant reduction in the amount of FLI generated in exposed rats compared to naïve rats. * indicate significant group differences (*p*<0.05).

**Figure 4:**
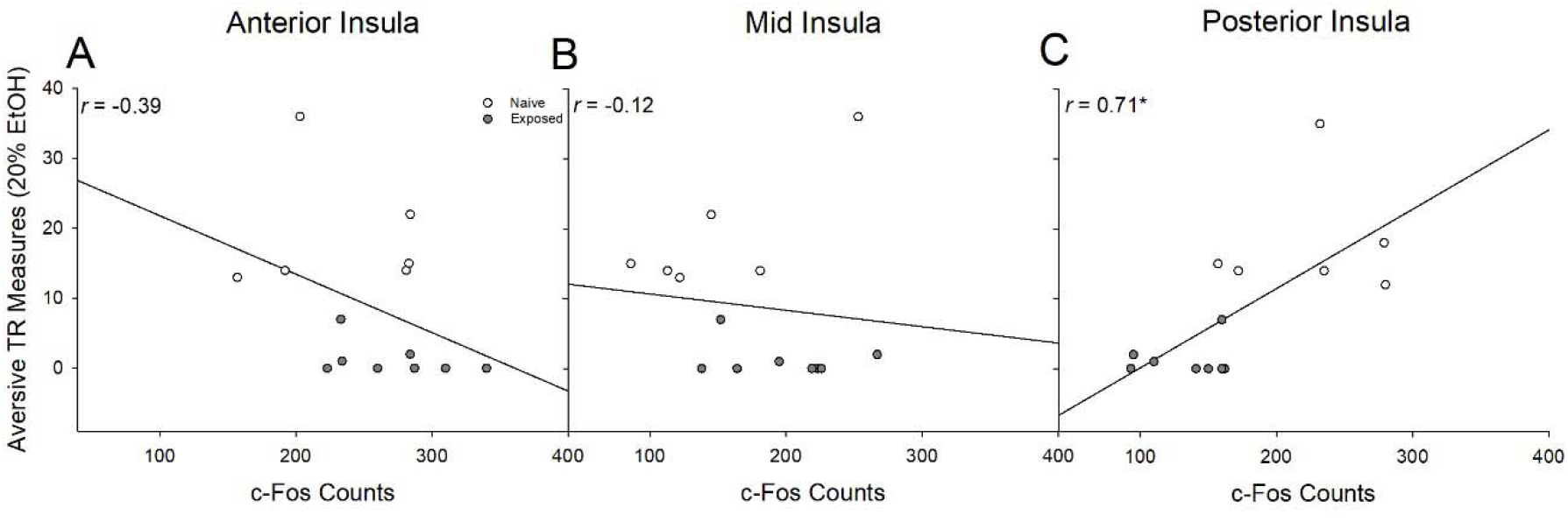
Changes in the aversive taste reactivity responses to orally delivered 20% EtOH were correlated with c-*Fos* expression in the insular cortex. **(A)** There was a strong negative correlation between the number of FLI+ cells in the anterior IC and aversive TR (r=0.39) which did not reach statistical significance. **(B)** No measurable correlation was observed between FLI+ cells in the middle IC and aversive TR (r=0.12). **(C)** A significant negative correlation was observed between the number of FLI+ cells in the posterior IC and aversive TR (r=0.71). * indicate significant group differences (*p*<0.05).

### 3.2 Inhibition of IC increases acceptance of higher concentrations of EtOH in naïve rats

Inhibition of IC induced a modest but statistically reliable increase in the acceptance of higher concentrations of EtOH in naïve-rats, and removal of IC inhibition eliminated these differences. A three-factor mixed ANOVA revealed a significant Test x Drug interaction (*F*_(1,17)_=7.83, *p*<0.05). When intra-IC infusions were given during the first three exposures to EtOH (Fig 5A), rats receiving GABA agonists licked more to EtOH relative to rats receiving PBS. On the following three test sessions, in which rats underwent sham infusions (Fig 5B), this difference was lost. Planned comparisons comparing licking at the highest concentrations of EtOH revealed that IC-inhibited rats licked significantly more to the 20 and 40% concentrations, relative to controls (Fig 5). This did not result in differential total g/kg EtOH consumed during these 30-min tests (0.50 ± 0.05 vs 0.55 ±0.08 g/kg/30min, respectively). Figure 6A is a representative image of a thionin stained coronal section of the IC taken to verify cannula placements for all animals that took part in this study and Fig 6B demonstrates the relative location of all injection sites within the IC.

**Figure 5:**
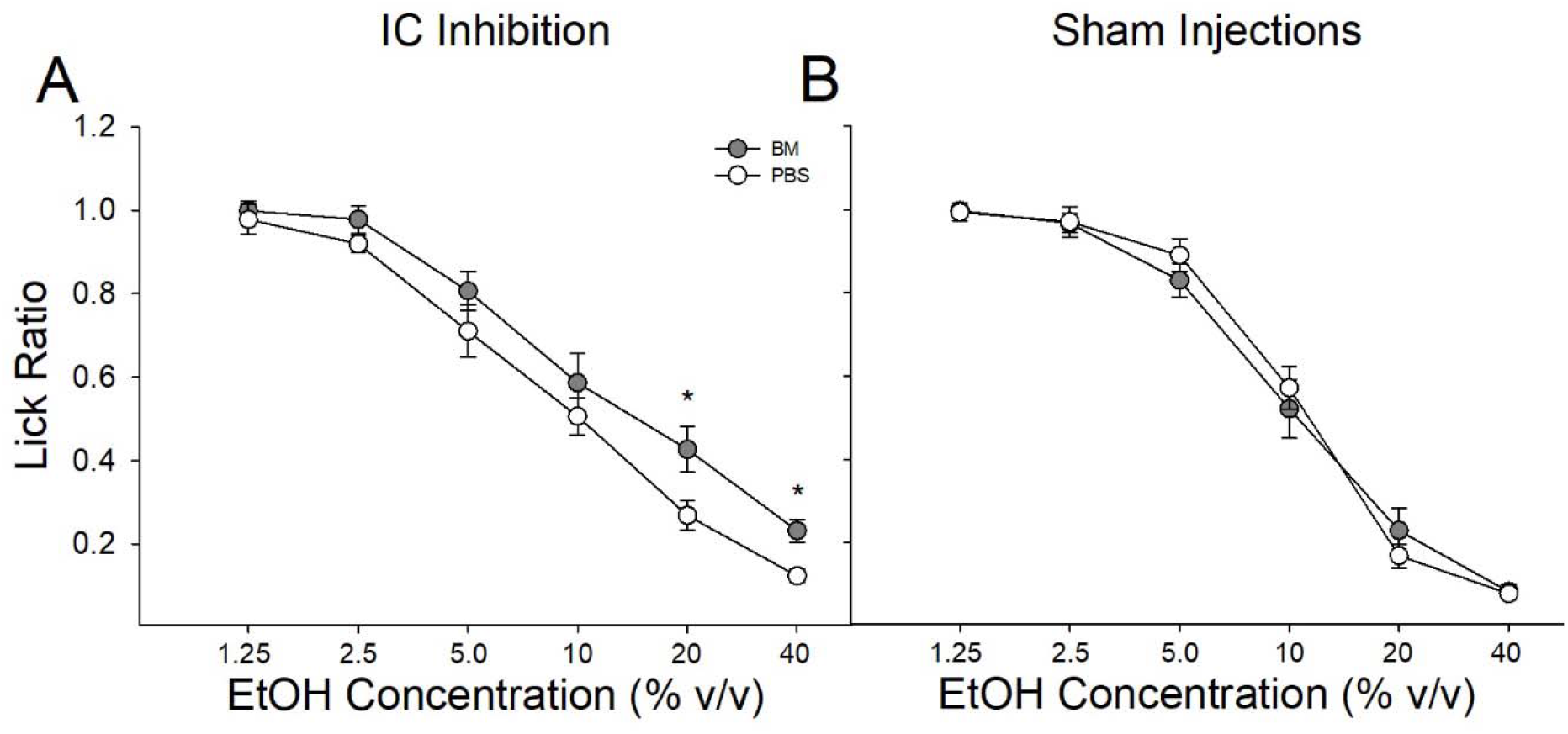
Pharmacological inhibition of the insular cortex (IC) prior to BAEE increases the acceptance of higher EtOH concentrations in naïve rats. **(A)** Naïve rats’ average licks to each EtOH concentration following the GABA agonist (baclofen-muscimol; BM) treatment or vehicle (PBS) into the IC. Inhibition of the IC resulted in significantly more licks at 20% and 40% EtOH in BM rats compared to PBS controls and **(B)** sham injections resulted in the collapse of this behavior. * indicate significant group differences (*p*<0.05).

**Figure 6:**
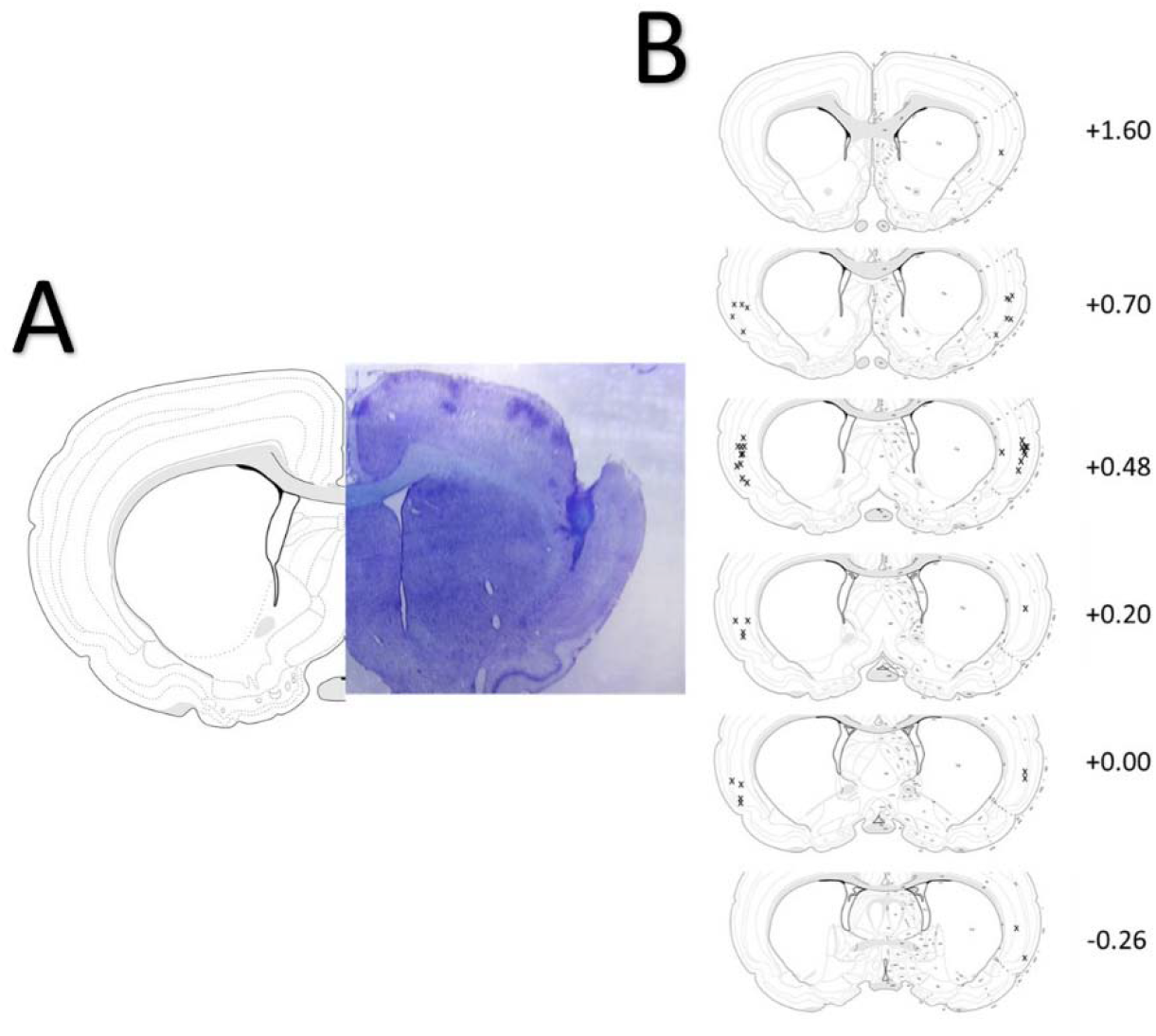
Cannula verification within the insular cortex (IC). **(A)** Representative thionin-stained image showing the tissue damage resulting from the cannula track targeting the IC. **(B)** Coronal sections showing approximate sites of the cannula tracks from bregma from all the rats included in Experiment 2.

#### 3.3.1 Escalation of EtOH intake within the BAEE model does not alter sensitivity to EtOH taste

BAEE-induced escalation of EtOH intake failed to alter sensitivity to the taste of EtOH as assessed with a psychophysical taste detection paradigm. A two-factor mixed ANOVA conducted on licking responses in the BAEE paradigm revealed a significant Test x Concentration interaction (Fig7A; *F*_(5,25)_ = 2.84, *p* < 0.05), demonstrating robust escalation of EtOH intake. We next compared naïve- and exposed-rats on their ability to report two concentrations of sucrose or QHCl as discriminable from H_2_O in two separate three-factor ANOVAs. For sucrose, this analysis revealed a Stimulus x Concentration (*F*_(1,10)_ = 9.88, *p* < 0.05) interaction such that rats performed better on the higher concentration of sucrose (600mM) relative to the lower concentration (200mM). For QHCl, we similarly found a Stimulus x Concentration interaction (*F*_(1,10)_ = 17.1, *p* < 0.01), again driven by the finding that all rats performed better on the higher concentration of QHCl (1.0 mM) compared to the lower concentration (0.4mM). Importantly, there were no effects of EtOH exposure on performance during these final stages of training, indicating that prior EtOH experience did not affect the ability to perform this task.

**Figure 7:**
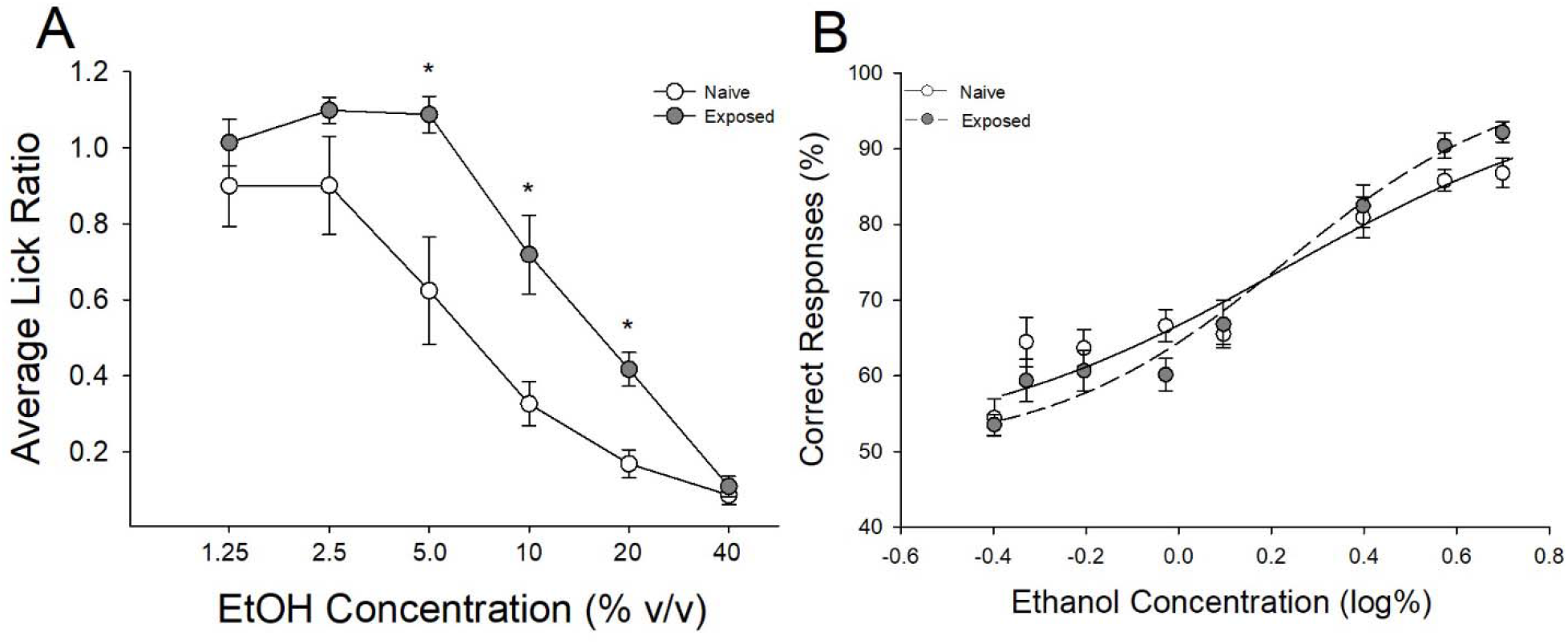
Escalation of EtOH intake following the BAEE paradigm does not alter sensitivity to the taste of EtOH. **(A)** Comparison of average lick ratios following initial exposure to EtOH (Naïve) and following six sessions of BAEE (Exposed). Rats show significantly more licks to 5%, 10% and 20% EtOH following six sessions of BAEE. **(B)** There were no significant differences in the ability to correctly identify the taste intensity of EtOH between Naïve and Exposed groups at any given concentration (displayed in log%). Data are expressed as the percentage of correct responses when asked to identify EtOH or water. * indicate significant group differences (*p*<0.05).

A two-factor ANOVA conducted on the proportion of correct responses at each of the eight EtOH concentrations revealed a significant main effect of concentration (Fig 7B; *F*_(7,70)_ = 81.64, *p* < 0.0001) such that performance dropped as a function of decreasing EtOH concentration. Importantly, there were no main or interactive effects of EtOH exposure on the proportion of correct responses. We next compared each of the curve parameters between exposed and naïve rats with a series of one-way ANOVAs. We failed to find an effect of EtOH exposure on either maximal responding (*a*), curve slope (*b*), or the concentration of half-maximal performance (*c*).

#### 3.3.2 Escalation of EtOH intake within the BAEE model does not alter the salient taste quality of EtOH

BAEE-induced escalation of EtOH intake did not alter the salient taste quality of EtOH regarding the degree to which the taste of EtOH generalized to the taste of either sucrose or QHCl. A two-factor mixed ANOVA on licking during the BAEE paradigm revealed a significant escalation of EtOH intake resulting in a significant Test x Concentration interaction (*F*_(5,40)_ = 2.85, *p* < 0.05). We first conducted a three-factor mixed ANOVA on the performance of EtOH-naïve and - exposed rats in being able to correctly identify the stimulus when presented with the sucrose and QHCl standards during testing. This analysis revealed a significant main effect of Stimulus (*F*_(1,15)_ = 16.29, *p* < 0.01), such that all rats were better at identifying QHCl within this paradigm relative to sucrose. Importantly, there were no main or interactive effects of EtOH Exposure, again indicating that prior EtOH experience did not affect the ability to perform this task. A two-factor mixed ANOVA conducted on Sucrose Generalization Scores between naïve and exposed rats as a function of EtOH concentration revealed a main effect of EtOH Concentration (*F*_(6,90)_ = 69.31, *p* < 0.0001), driven by rats increasing their responding on the sucrose-associated side as a function of increasing EtOH concentration. Importantly, there were no effects of EtOH Exposure on the Sucrose Generalization Scores, indicating that EtOH exposure did not significantly impact the degree to which the taste quality of EtOH resembled sucrose or QHCl.

## 4.0 Discussion

We examined the parameters associated with rapid escalation of EtOH intake that is observed following the BAEE paradigm. In Experiment 1, we determined the effect of BAEE-induced escalation of EtOH on subsequent changes in hedonic responding to orally delivered 20% EtOH. Subsequently, we examined neural response within the IC following orally-delivered EtOH as approximated through quantification of FLI+ neurons. First, we replicated previous findings that 6 non-consecutive sessions consisting of intermittent brief-access ethanol exposures significantly increased oral self-administration of EtOH and increased the acceptance of higher concentrations of EtOH (i.e. ≥5.0%) as determined by a significantly enhanced lick rate (Fig 1A & B). It is important to note that rats are only required to make one lick to a given concentration of EtOH before being presented with water and then a different EtOH concentration. This ensures that all animals sample each EtOH concentration but are not required to measurably consume any given concentration. Our finding that rats voluntarily increased lick rate to these higher concentrations is demonstrative of a substantive change in the evaluation of concentrated EtOH. Consistent with this interpretation, EtOH-exposed rats displayed significantly reduced aversive oromotor taste reactivity (Berridge, 2000, Grill and Norgren, 1978, Grill and Berridge, 1985) responses to intraorally delivered 20% EtOH, relative to naïve rats, indicating that the aversive properties of EtOH were specifically impacted following EtOH escalation (Fig 2). It was previously demonstrated that more prolonged exposure to EtOH produced a similar decrease in aversive responding to EtOH, albeit at a lower concentration (12% vs 20%; Kiefer and Dopp, 1989)

Subsequent quantification of FLI+ neurons across three anterior-posterior sections of IC revealed alterations in neural responsivity to intraoral 20% EtOH, which agrees with these behavioral effects. EtOH-exposed rats displayed a reduced expression of FLI+ cells in posterior IC following delivery of EtOH (Fig 3). The number of FLI+ cells in the anterior and mid sections of IC did not differ between EtOH-exposed and -naïve rats. Next, we conducted a series of regression analyses to correlate the expression of FLI+ cells within the three sections of the IC with the number of aversive taste reactivity measures (Fig 4). We found a significant positive correlation between the number of FLI+ cells within the posterior IC and the number of aversive oromotor responses. This finding is consistent with previous work demonstrating a positive correlation between the number of *Fos*+ neurons in a similar posterior subregion of IC and the number of aversive taste responses produced in rats receiving intraoral infusions of innately aversive QHCl solutions (King, 2018). Opioid signaling within this area of IC increases the positively hedonic “liking” response elicited by intraorally delivered sucrose solutions (Castro and Berridge, 2017) further demonstrating that neural adaptations within this subregion of IC may be critical for driving enhancements of hedonic responding to orally delivered stimuli. Furthermore, deletion of dynorphin within IC blocks escalated EtOH intake in mice (Pina et al., 2021). Interestingly, the number of aversive responses to EtOH was negatively correlated with the number of FLI+ cells within the anterior IC, although this difference did not quite reach statistical significance. Increased anterior IC activity is implicated in the expression of compulsive-like EtOH intake that is insensitive to the aversive outcomes of EtOH consumption (Campbell et al., 2019, Darevsky and Hopf, 2020). Altogether, these data strongly suggest that IC, particularly the posterior IC, may be critically responsible for the escalation of EtOH intake and associated concurrent decrease in aversive responding to EtOH.

In an attempt to directly demonstrate this relationship between IC and voluntary EtOH intake at higher concentrations, we next sought to inhibit IC in naïve rats and examine EtOH consumption in the BAEE paradigm. We found that acute pharmacological inhibition of IC was sufficient to modestly, but statistically reliably, augment initial acceptance of higher EtOH concentrations in naïve-rats. Specifically, pharmacological inhibition of IC in EtOH-naïve rats resulted in increased licking at higher EtOH concentrations (20 & 40%) compared to PBS controls (Fig 5A), and this difference was lost when the pharmacological treatments were removed (Fig 5B). It is important to note that these differences in licking did not translate to increased consumption (g/kg) of EtOH. One potentially important caveat to this study is that the IC cannulae were implanted just anterior to the area of the IC in which we observed reduced FLI+ neurons following EtOH escalation (Fig 6). It is possible that, had these cannulae been positioned more posteriorly, we may have observed a larger effect of IC inhibition on EtOH consumption. These studies are currently planned as follow-up experiments.

Given the findings of our lab, and others, that IC is critically involved in both escalated and compulsive-like EtOH consumption (Centanni et al., 2019, Seif et al., 2013, Chen and Lasek, 2020) we next sought to definitively determine the degree to which taste perception of EtOH changes following sufficient experience to drive escalated EtOH consumption. To this end, we employed two psychophysical paradigms designed to determine the degree to which EtOH-naïve and -exposed rats differed in responding to the taste properties of EtOH. There are three key perceptual domains along which taste stimuli can vary: hedonics, intensity, and quality (Spector, 2000). Data from the BAEE paradigm (Fig 1) and taste reactivity measures (Fig 2) conclusively demonstrate that EtOH-exposed rats differ from naïve rats in their hedonic responding to EtOH, so we designed two experiments to independently test for differences in the intensity and, separately, the salient taste quality of EtOH as a function of prior EtOH exposure. Initially, one group of rats was put through the BAEE paradigm to escalate EtOH consumption while the other group remained naïve (Fig 7A). Then, we employed an adaptation of a taste detection paradigm to examine the degree to which rats could successfully identify the taste of EtOH as concentration was systematically lowered. We found that EtOH-naïve and - exposed rats did not differ on any measure of sensitivity to the taste of EtOH as assessed with this paradigm (Fig 7B). All rats were capable of competently performing the task, and all rats were similarly impacted in their performance as the EtOH concentration was decreased. Thus, EtOH exposure did not appear to affect the perceived intensity of the taste of ethanol as assessed within this paradigm.

Prior findings demonstrate that oral EtOH activates both sucrose and QHCl responsive taste pathways and that the taste quality of EtOH is often generalized to ‘bitter-sweet’ taste mixtures (Lanier et al., 2005, Scinska et al., 2000, Bartoshuk et al., 1994, Kiefer et al., 1990, Di Lorenzo et al., 1986, Nolden et al., 2016, Nolden and Hayes, 2015, Mattes and DiMeglio, 2001, Green, 1988, Lemon et al., 2004, Kiefer and Mahadevan, 1993). Exposure-induced changes in the salience of either taste properties could significantly impact the hedonic responding to EtOH taste. As such, in a separate group of rats, we tasked animals with reporting the degree to which the taste of EtOH resembled that of either sucrose or QHCl as a function of prior EtOH exposure. Again, rats either remained naïve or were exposed to EtOH in the BAEE paradigm (Fig 8A). Following exposure, we examined the degree to which rats generalized the taste of EtOH to sucrose or QHCl in an operant task. We found that all rats, regardless of EtOH exposure history, reported the taste quality of EtOH became more sucrose-like as the concentration of EtOH increased. Importantly, at no point did EtOH-naïve and -exposed rats differ in their treatment of the taste of EtOH with regard to how perceptually similar it was to sucrose or QHCl. Together, these two experiments demonstrate that escalation of EtOH intake within the BAEE paradigm is not representative of sizable alterations in the perceived taste properties of EtOH in rats. Rather, the driving phenomenon appears to be hedonic or motivational in nature. Prior research has demonstrated that rats that displayed an EtOH-exposure induced increase in acceptance of concentrated EtOH solutions also displayed significant reductions in the activity of taste-nerve fibers in response to orally-delivered EtOH, albeit in adolescent rats and following a much more protracted EtOH exposure period (Tang et al., 2018). While our data demonstrate no change in the psychophysical assessment of the taste of EtOH following escalated EtOH intake in adult rats it is important to address the caveat that a more extensive EtOH exposure period or greater total EtOH intake may potentially alter the taste properties of EtOH. Rather, our present data strongly indicate that changes in the taste properties of EtOH are not required for escalation of EtOH intake.

**Figure 8:**
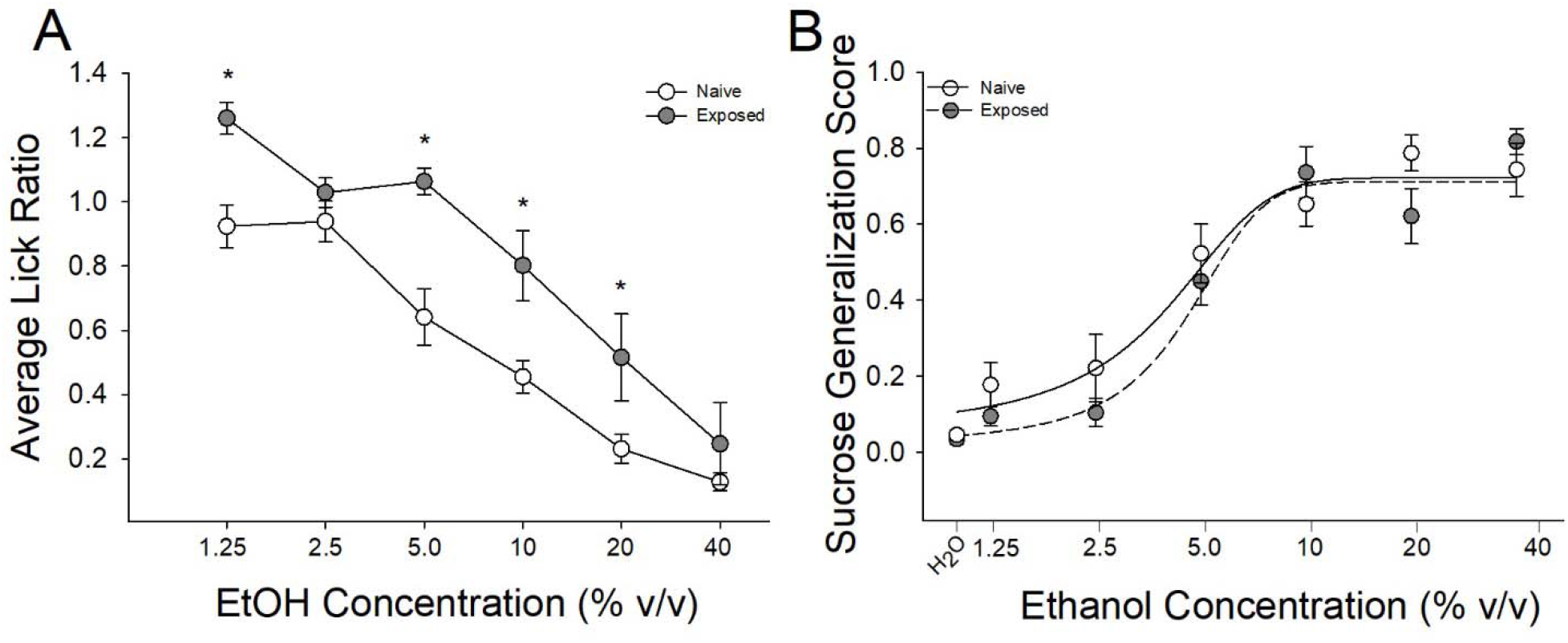
Prior escalation of EtOH intake following the BAEE paradigm does not alter the salient taste quality of EtOH. **(A)** Comparison of average lick ratios following initial exposure to EtOH (Naïve) and following six sessions of BAEE (Exposed). Rats show significantly more licks to 1.25%, 5%, 10% and 20% EtOH following six sessions of BAEE. **(B)** As the concentration of EtOH was increased, naïve and exposed rats reported that the taste of EtOH generalized more to sucrose rather than QHCl. A Sucrose Generalization Score of 1 means that the rat responded consistently on the sucrose associated side and a Sucrose Generalization Score of 0 means the rat responded consistently on the QHCl associated side. There were no significant differences in Sucrose Generalization Score between naïve and exposed rats at any given EtOH concentration. * indicate significant group differences (*p*<0.05).

In summary, we have confirmed that just six BAEE sessions is sufficient to drive escalated EtOH intake consistent with our previous report (Loney and Meyer, 2018). Furthermore, this escalation is associated with concomitant changes in IC neural activity and a decrease in sensitivity to the aversive components of EtOH and this change is associated with altered IC neural activity in a manner that does not appear to affect the salient taste properties of EtOH. These data further implicate the IC as an important neural locus maintaining control over problematic EtOH consumption. Future studies should address the relevant corticofugal and corticocortical connections of the IC in attempt to restore sensitivity to the adverse consequences of EtOH consumption.

## Acknowledgments

We thank Kelechi Ezenwa, Zoe Giandomenico, Mackenzie Haller, Renee Rouleau, and Max Towers for their valuable contributions. We would also like to thank Dr. Ann-Marie Torregrossa’s lab for the use of testing equipment. The authors have no conflicts of interest to declare.

