## Supplementary for "Escalation of alcohol intake is associated with regionally decreased insular cortex activity but not associated with changes in taste quality"

The following are detailed versions of the Davis rig and gustometer methods which were abbreviated in the manuscript due to word limits imposed by the journal:

**2.4.2 Brief Access Ethanol Exposure (BAEE)**

*Training.* Prior to training, animals underwent overnight water deprivation. All training sessions lasted for 30 minutes and consisted of tap water as the sole stimulus. On the first training day, animals were provided access to one stationary sipper tube of water and allowed to freely lick for 30 minutes. On the second day, animals received seven sipper tubes of water and were habituated to the movement of the shutter and the sliding table carrying the sipper tubes. During this training session, rats were given 60 seconds to initiate a lick response. Once a lick was counted, rats were allowed 30 subsequent seconds to freely lick. This 30-s period was followed by a 10-s intertrial interval prior to commencing a new trial. A new trial was presented to the animal even if the rat failed to initiate a lick on the current trial. On the third day, rats were given 30 second presentations followed by a 1-s presentation of a subsequent water spout with a 10-s intertrial interval. A new trial was not presented in the case that an animal failed to lick at any presentation. On the fourth and final day of training, the access period to sipper tubes was shortened to 10 seconds. Each trial was followed by a 10-s intertrial interval followed by presentation of a new water spout for 1 second, which would serve as a water rinse during testing. Following this rinse, there was another 10 second inter-trial interval prior to commencing a new trial. Subsequent trials were not presented if the rat failed to lick during the current trial. This design ensures the animal is actively sampling each concentration of the stimuli. Importantly, the animal only has to lick one time to a given trial before being presented with a new trial and therefore is not forced to consume any concentration of any stimulus to appreciable levels.

*Testing.* Test sessions were identical to the final day of training except that animals were presented with a randomized block consisting of six concentrations of EtOH (1.25, 2.5, 5.0, 10, 20 & 40%) and water. Licks on each concentration were recorded, and animals were able to initiate as many trials as possible during the 30-minute session. Testing was conducted intermittently on Monday, Wednesday, and Friday of a given week, allowing approximately 48 hours in between test sessions. Immediately following testing, the animals received *ad libitum* food and water overnight in their home cage.

**2.6 Experiment 3a: Ethanol Taste Detection Threshold Assessment in Naïve and Exposed Rats under the BAEE Model**

**2.6.1 Animals**

Male and female rats were weight-matched into two groups: exposed (n=6; 3 males, 3 females) and naïve (n=6; 3 males, 3 females). On all weekdays, animals were maintained on a 23-hour water deprivation schedule, such that they would be motivated to earn water during the daily 30-minute sessions in the gustometer. Water was removed ~24h before the first session and returned after completion of the last session of a given week, and water was available *ad libitum* on weekends. Animals were weighed and handled daily, and standard chow was provided *ad libitum* throughout the duration of the experiment.

**2.6.2 Brief Access Ethanol Exposure (BAEE)**

Training and testing procedures were identical to experiments 1 & 2. Water was removed prior to BAEE sessions. After training, rats were split into two groups for testing: an “exposed” group that would be given access to various concentrations of EtOH (1.25, 2.5, 5.0, 10, 20 & 40%) and water, while a “naïve” group would only be presented with water during testing. Licks on the sipper tubes for each concentration were recorded through the lickometer system.

**2.6.3 Gustometer Spout Training**

Rats were initially trained to lick a dry sample stimulus delivery ball located in the center of the front of the test chamber. When the rat initiated at least two licks with an inter-lick interval (ILI) of ≤ 250 ms, a 5 µl aliquot of water from one of several fluid reservoirs was deposited onto the sample ball. Subsequently, rats could elicit up to five more licks (∼5 μl of fluid per lick) during the 3 s sample period before the sample ball was removed and cleaned with water. On subsequent days, each rat underwent identical training sessions for the right side and left side spouts, also located through slots in the test chamber side wall as described above. This served to familiarize rats with the device and trained them to obtain fluid from each ball. All rats had access to the same stimulus ball on any given day, and training lasted for 3 days, 1 day per ball. The duration of all training and testing sessions was 30 min.

**2.6.4 Gustometer Side Training**

After spout training, rats were conditioned to associate a given reinforcement spout with either the standard taste stimuli (sucrose and QHCl) or water. Side assignments were counterbalanced across animals. Immediately after stimulus sampling, rats were given a 180 s decision phase (limited hold) to make a response on the correct assigned reinforcement spout. To facilitate the association, access to the unused reinforcement ball was occluded for that particular training day. Correct responses that occurred within the limited hold were reinforced with up to 20 licks of water (or 10 s access, whichever came first). Reinforcement volumes and duration were kept constant across all stages of training and testing. Only one standard stimulus (sucrose, QHCl, or DI water) was presented per day, and only the highest concentration of each training compound was used (600 mM sucrose and 1.0 mM QHCl, respectively). The remaining concentrations of standard stimuli were introduced on alternating days with a taste or non-taste stimulus. Side training lasted a total of eight days of training, consisting of two iterations of the following sequence: 1.0 mM quinine on D1, water on D2, 600 mM sucrose on D3, water on D4. This resulted in two stimulus-side pairings for each stimulus.

**2.6.5 Gustometer Alternation Training**

At this stage, rats were presented with one of two taste stimuli (1.0 mM QHCl or 600 mM sucrose) repeatedly until a criterion correct number of responses on the associated reinforcement ball occurred (6, 4, or 2 on days 1, 2, and 3 of training with a given stimulus, respectively). Correct responses did not have to occur sequentially. Once the criterion for that given stimulus was met, the rat was presented with water and required to reach the same criterion of correct responses on the associated reinforcement ball before the original stimulus was returned. This pattern was repeated throughout the session. During alternation training, the limited hold was reduced from 180 to 15 s, and a 10 s timeout was delivered in the case of an incorrect response or response omission. In an attempt to encourage generalization of the left side spout to all tastant stimuli, regardless of taste quality, we initially trained rats by presenting 1.0 mM QHCl, 600 mM sucrose, and water all within a single session, but rats struggled to demonstrate improvement on the task after repeated sessions, particularly for QHCl. As such, we lessened the task difficulty at this stage by giving rats alternating days on 1.0 mM QHCl and DI water versus 600 mM sucrose and DI water.

**2.6.6 Gustometer Discrimination Training Stage (I–II)**

For Stage I of discrimination training, we lowered the concentration of each standard stimulus (200 mM sucrose and 0.4 mM quinine). As above, only one standard taste stimulus was tested on a given day, and testing between this stimulus and water occurred across randomized trials. The limited hold to respond remained at 15 s after sampling, and the timeout for an incorrect response or omission remained at 10 s. After reaching a set criterion (~80% correct discrimination of tastant from water trials), animals moved on to Stage II. During stage II of discrimination training, two concentrations of quinine (1.0 & 0.4 mM) or sucrose (600 & 200 mM) were presented in randomized trials with water and only one taste stimulus was tested per day. Here the timeout for incorrect responses or omissions was increased to 20 s. On this final stage of training, it was found that EtOH naïve and exposed rats were performing substantially worse on 0.4 mM QHCl (57.9 ± 6.0 and 58.0 ± 5.8% correct, respectively) relative to 1.0 mM QHCl (89.8 ± 5.4 and 87.5 ± 5.6%). As such, a decision was made to give all animals five more days of training between H_2_O and 0.4 mM QHCl. At the end of this period, performance on 0.4 mM QHCl increased to 87.2 ± 4.3 and 85.0 ± 3.0% correct for EtOH naïve and exposed rats, respectively, so all rats were advanced to the testing phase of the experiment

**2.6.7 Testing EtOH Detection Threshold**

The testing phase was modeled after Stage II discrimination training, with the exception that only one EtOH concentration was presented per session (5%, 3.75%, 2.5%, 1.25%, 0.94%, 0.63%, 0.47, 0.4% v/v EtOH) along with H_2_O. Each concentration of EtOH was tested on two separate days, and performance was averaged across these days. In order to maintain stimulus control, each test day in which a lower concentration of EtOH was presented was followed with a test day in which rats were presented with the highest concentration at which they could reliably perform at, or above, 80% correct. This stimulus control day consisted of testing with 5% EtOH until it was determined that rats performed at, or above, 80% correct on 3.75% EtOH, at which point 3.75% became the concentration used on stimulus control days. At the end of testing, rats were given two test days on which all fluid reservoirs were filled with water instead of EtOH and water. This was done to ensure that animals performed at chance levels when the taste of EtOH could not be used as a cue, indicating that they were not using extraneous cues to guide responding.

**2.7 Experiment 3b: Assessment of the Salient Taste Quality of Various Ethanol Concentrations in Naïve and Exposed Rats Under the BAEE Model**

**2.7.1. Animals**

Male and female rats were counter-balanced based on weight and then assigned to either an EtOH-exposed group (n=9; 5 females, 4 males) or a naïve group (n=8; 4 females, 4 males). On all weekdays, animals were maintained on an ~23-hour water deprivation schedule, such that they would be motivated to respond for water in the gustometer during the daily 30-minute sessions. Water was provided *ad libitum* on weekends. Animals were weighed and handled daily, and standard chow was provided *ad libitum* throughout the duration of the experiment. These animals were experimentally naïve but had previously undergone a sham intercranial infusion surgery four weeks prior to training.

**2.7.2 Brief Access Ethanol Exposure (BAEE)**

Rats underwent identical EtOH exposure training and testing as previously described above. For the testing phase, naïve animals received water from all spouts during Davis rig sessions, whereas exposed rats were given 30-min access to six concentrations of EtOH (1.25, 2.5, 5, 10, 20, & 40% EtOH v/v) or water in a randomized fashion in the Davis rig. Licks on the sipper tubes for each concentration were recorded through the lickometer system.

**2.7.3. Gustometer Spout Training**

As in Experiment 3a, rats were given a three-day training that required learning to lick a dry sample stimulus delivery ball at the center, right, or left in order to elicit water delivery. Training lasted for 3 days, 1 day per spout. The duration of all training and testing sessions was 30 minutes.

**2.7.4. Gustometer Side Training**

Rats were trained to associate a given reinforcement ball with a standard stimulus (either sucrose or QHCl). Rats were presented with only one standard stimulus per day, and, during this phase of training, only the highest concentration of each stimulus was used (600 mM sucrose or 1.0 mM QHCl). All other conditions for side training were identical to those described in Experiment 3a. As such, the standard stimulus used during training was alternated daily for every rat, and animals were tested on the subsequent day with the other stimulus. All rats were presented with a 180 s limited hold after sampling the stimulus in which to make a response on the stimulus-associated reinforcement ball. The unused balls were covered and thus not available to the rat for that particular side training day. If the rat responded correctly, their response was reinforced with 20 licks (or 10 s access, whichever came first) of water. Reinforcement volumes and duration were kept constant across each stage of training and testing. Regardless of outcome (response or no response), there was a 6 s intertrial interval. Side training lasted for four days, with two days of training allotted for each stimulus/spout assignment.

**2.7.5. Gustometer Alternation Training**

During this phase of training, both the left and right reinforcement spouts were made available. Alternation training sessions consisted of presenting one of the two standard stimuli (600 mM sucrose or 1.0 mM quinine) repeatedly until the rat reached a criterion number of correct responses (6, 4, or 2 on training Days 1, 2, and 3, respectively). These correct responses were not required to be sequential. Once criterion was reached, the other stimulus was presented and the animal was required to reach the same correct response criterion before returning to the original stimulus. This pattern was repeated for the entire duration of the 30-min test. The limited hold for alternation training was reduced from 180 to 15 s, and a 10 s timeout was delivered in the case when a rat made an incorrect response or failed to make any response.

**2.7.6. Gustometer Discrimination Training (Stage I-II)**

Discrimination training was completed across three stages. For the first stage of discrimination training, only the intermediate concentrations of the standard stimuli (200 mM sucrose and 0.3 mM QHCl) were presented in randomized blocks. Rats were tasked with making a choice regarding the taste quality of the sample stimulus by responding on one of the two reinforcement spouts. The limited hold was maintained at 15 s, and incorrect or omitted responses were punished with a 10 s timeout. After reaching a performance criterion (~80% correct for both stimuli), animals were advanced to Stage II of discrimination training, which consisted of presenting all three concentrations of both sucrose and QHCl presented in blocks of six randomized trials (without replacement). For sucrose, the three concentrations provided were 60, 200, and 600 mM, and, for quinine, the concentrations were 0.1, 0.3, and 1 mM. For Stage II, the timeout for incorrect or omitted responses was increased to 20 s. Finally, Stage II discrimination training parameters served as a training refresher on the Monday sessions during testing.

**2.7.7. Taste Discrimination Testing**

The testing phase modeled Stage II discrimination training, although the varying EtOH concentrations were introduced on Tuesday–Friday of each week. Monday of any given week served as a stimulus control day. in which only sucrose and QHCl were presented, with concentrations and randomized presentation of stimuli identical to Stage II discrimination training parameters. Two concentrations of EtOH were tested within a given week, and each concentration of EtOH was tested twice, with performance averaged across the two testing days. The concentrations of EtOH tested consisted of the same concentrations used in the Davis rig (1.25, 2.5, 5.0, 10, 20, 40% v/v). As above, to ensure that animals were responding solely to the chemical properties of the training stimuli, two water control sessions were conducted in which each reservoir was filled with DI water and arbitrarily assigned as one of the three concentrations of each standard stimulus.
